# Improving plant functional annotation from knowledge graphs using Graph Neural Networks

**DOI:** 10.1101/2025.05.07.652750

**Authors:** Tran Gia Bao Ngo, Christophe Liseron-Monfils, Shreyan Das, Jordan Ubbens, Paula Ashe, David Konkin

## Abstract

Annotating genes is essential to crop development, and understanding gene functions sheds light on developing crop improvement strategies, such as marker-assisted breeding, genetic modification, or pest resistance. Through an extensive experimental effort and computational annotation projection, tens of thousands of genes have been annotated across plant species, with most of the gene annotations focusing on a well-studied species, Arabidopsis thaliana, but this represents a small fraction of the hundreds of thousands of genes across these different plant species. Phenotypes and their traits result from multiple processes and events involving multiscale information encoded from different omics, such as genomes, proteomes, or transcriptomes. This stresses a need for an efficient computational approach to capture and integrate information from biological networks and transfer this knowledge from well-studied species to unknown species to annotate and discover functional relationships between annotations and genes. Despite recent progress, existing methods only consider one or a few omics levels to perform reasoning on functional annotation-to-gene relations. The main objective of this study is to generate and explore a large-scale plant biological knowledge graph, the DasDB, and to enrich gene functional annotation linked to genes in different species using graph neural networks (GNNs). Integrating various data sources from different omics has resulted in a comprehensive graph database, facilitating researchers’ in-depth understanding of complex biological networks at the highest level. In addition, applying GNNs on a large-scale knowledge graph database has shown promise in the ability of deep learning models to transfer this information from well-studied plant species to less-characterized plant species, outperforming the transfer of information done using only orthology relationships. This study benchmarks a new research direction in producing new functional annotation discovery in plant species with limited functional annotations. This pipeline was applied to a specific research problem: the mechanism involved in pea nodule nitrogen fixation. We identified known gene markers of this process through a systematic analysis of the DasDB, showing the relevance of our approach. Furthermore, new potential targets to better understand and improve this process were identified.

## 1 Introduction

The emergence of high-throughput biotechnology has increased the availability of diverse omics data, including genomics, epigenomics, transcriptomics, proteomics, and metabolomics, which complement each other to represent complex biological networks. Recent advances in machine learning and deep learning have helped obtain new insight from these data to produce classification biomarkers. New biomarker generation approaches only consider single-omic data, which lacks the precision to establish robust associations between molecular-level changes and phenotypic traits (Valous et al., 2024). Multi-omic data combines complementary knowledge from across different omics layers, enabling deep neural networks to effectively discover hypothesis-generating biomarkers in many biological applications, such as biomedicine (Xiao et al., 2023; Hasin et al., 2017; Gong et al., 2023), cancer biology (Menyhárt and Győrffy, 2021; Cai et al., 2022; Poirion et al., 2021; Akhoundova and Rubin, 2022; Schulte-Sasse et al., 2021; Li et al., 2022; Gao et al., 2022), and crop genomics. However, these data types are unavailable for studies of legumes such as peas. A possible way to gain information for a species such as pea (*Pisum sativum*) is to leverage the knowledge created in other plant species, such as *Arabidopsis thaliana*. An effective technique to achieve this knowledge transfer is to generate knowledge graphs. In plant research, graph knowledge has been developed (Hassani-Pak et al., 2021; Confais and Francillonne, 2023; Larmande and Todorov, 2021; Zhang et al., 2024). These resources are excellent if only public datasets are required to help generate hypotheses by transferring knowledge between species. However, these public resources are more difficult to use for lab-specific datasets and can lack the flexibility to analyze lab-specific data on less-studied plant species. A recent, legume-centered, knowledge graph was established, using legume-specific data information. However, this resource considered the knowledge resources from *Arabidopsis thaliana* outside the legume knowledge spaces (Imbert et al., 2023). The increased availability of tools in this space is useful. However, researchers need to be able to create knowledge graphs that they can adapt to their tailored research questions.

Machine learning or biostatistical methods analyze multi-omic data from different sources with more or less robust outcomes (Chicco et al., 2023). Due to their expressive representation capabilities, deep neural networks (DNNs) have been adopted, along with the advantages of multi-omics data, to analyze and capture patterns hidden behind complex biological networks. DNN-based approaches can be summarized in three stages: (a) Transforming the high-dimensional features into high-level semantic embeddings; (b) Learning a unified representation from the multiple embeddings; and (c) Applying the learned representation to downstream tasks (Xiao et al., 2023). Graph structure-based data, such as protein-protein or gene-gene interactions, has existed for decades in biology prior to the introduction of biological knowledge graphs. The analysis of these graph-based data is complex for traditional deep learning methods based on Euclidean data, having a finite space representation (Khemani et al., 2024; Sapoval et al., 2022). A different strategy for omics datasets is to model them as graph-structured data so that the relevant entities can be connected based on their intrinsic relationships, biological properties/significance, and empirical biological knowledge. All interactions within and across different omics sets form an interlinked graph (network) composed of vertices (nodes or entities) and edges (links or relationships) (Valous et al., 2024). This graph-based representation of multi-omic data was efficiently modelled by Graph Neural Networks (GNNs), in various biological tasks (Xiao et al., 2023; Schulte-Sasse et al., 2021; Wang et al., 2021b; Li et al., 2022; Kim et al., 2015; Gao et al., 2022; Huang et al., 2024; Gysi et al., 2020). GNNs effectively learned a representation of the data embedded in graph structure, capturing relationships within and across omics layers (Valous et al., 2024). In recent years, GNNs have become powerful and functional tools for machine learning tasks in the graphical domain. This progress is due to advances in expressive power, model flexibility, and training algorithms (Zhou et al., 2020).

Despite recent advancements in GNNs and the success of multi-omic data in various biological domains, leveraging GNNs on multi-omic data on plant biological tasks to improve crops has yet to be explored, especially for annotating unknown genes in crops. Guilt-by-association-based methods (Wolfe et al., 2005; Jiao et al., 2017; Gillis and Pavlidis, 2011b) have been the primary concept to solve the gene functional annotation task. However, the dominant effect of genes with high connectivity within molecular networks potentially misleads the prediction of new annotation (Gillis and Pavlidis, 2011a). On the other hand, recent works (Sehrawat et al., 2023; Zeng et al., 2021; Ma et al., 2018) have adopted deep learning techniques to learn a representation of genes according to a single layer of omics data. This approach has been limited since phenotypes and their trait components result from multiple processes and events across numerous omics layers, such as the genome, proteome, transcriptome, epigenome, etc. Thus, the primary focus of this work is to address this gap by showcasing the advantages of multi-omic graph data combined with GNNs on gene functional annotation tasks. Assessment of the model functionality is critical to determine its suitability for detecting patterns in data, the biological relevance, and the accuracy of the prediction, by comparing them with known contributors to the studied biological function and metabolic pathways (Sapoval et al., 2022).

The utilization of synthetic nitrogen (N) fertilizer is an environmental problem due to its atmospheric-polluting production and its under-utilization by crops, which leads to water pollution. Legumes can transform atmospheric nitrogen to plant-available nitrogen thanks to a symbiotic interaction with bacteria in a specialized organ called nodules in the roots (Stougaard, 2000). Within nodules, the bacteria transform into a state called bacterioid, fixing nitrogen for the plant in exchange for sugar resources derived from the plant photosynthesis. This nitrogen fixation benefits the pea itself and intercropping, enabling reduced application of inorganic nitrogen fertilizer (Toker et al., 2024). Systemic nitrogen signaling modulates symbiotic nodule functions, notably N uptake in peas. Consequently, the nodule example to test the DasDB was performed in low nitrogen conditions to study the symbiosis effect on nitrogen-related genes in conditions that enable nodule nitrogen fixation capacity.

This study develops DasDB, a large-scale plant biological knowledge graph integrating multi-omics data to enhance gene functional annotation. By applying Graph Neural Networks (GNNs), it improves the transfer of functional annotations from well-studied to less-characterized plant species, outperforming traditional orthology-based methods. The approach was validated on pea nodule nitrogen fixation, identifying both known gene markers and new potential targets, showcasing its potential for broader crop improvement applications.

## 2 Materials and methods

To build the DasDB, a comprehensive multi-omic database revealing a complex picture of plant biology, we integrate various open-source databases, namely Ensembl Genomes, Plant Gramene, OBO Foundry, EBI Expression Atlas and Bio GRID, into a unified graph database. In Section 2.1, we provide detailed documentation on the data pre-processing and database construction pipeline. We then perform knowledge graph reasoning to predict associated functional annotations of genes by using a GNN. We choose the state-of-the-art A*Net (Zhu et al., 2022) as an example experiment architecture. Section 2.2 explains our choice and offers a brief overview of the study methodology.

### 2.1 DasDB: Multi-omic graph database

The heterogeneous graph representation of multi-omics data provides an advantage for discerning functional patterns for predictive/exploratory analysis, enabling the modelling of complex biological relationships (Valous et al., 2024). In this work, we build a comprehensive multi-omic graph database by integrating various data sources with an end-to-end pipeline described in Figure 1. The multi-omics knowledge graph was built using Neo4J version 4.4.31 (March 2024). We first fetch raw data from Ensembl Genomes for genomic data, Plant Gramene for pathway and metabolite data, OBO Foundry for ontology data, EBI Expression Atlas for transcriptomic data and Bio GRID for proteomic data. In addition, we augment the database with internal experimental data. The pathway data were obtained from the Plant Reactome Naithani et al. (2019). After fetching all raw data, we perform data standardization and create Snakemake pipelines to facilitate fetching and cleaning data (Mölder et al., 2021). We populate the Neo4J database using the cleaned data, then run OrthoFinder2 (Emms and Kelly, 2019) to find Orthogroups, which are added back to the database. Our multi-omic graph database represents a complex biological network with more than 16 million different omics nodes and 170 million relations between entities, comprised of 17 plant species and 12 omics layers. Details about the number of entities for each omic are shown in Table 1. To facilitate access to the data, a Graph API was generated to prepare possible data transfer to a user interface using the GraphQL API.

**Table 1:** Number of entities (nodes) for each data type in graph database.

| Data type | Number of entities |
| --- | --- |
| SNP | 12,894,865 |
| Functional Annotation | 2,620,206 |
| Species | 17 |
| Gene | 744,192 |
| mRNA | 185,921 |
| Protein | 26,620 |
| Orthogroup | 43,498 |
| Omics Sample | 15,792 |
| Pathway | 15,708 |
| Reaction | 11,868 |
| Metabolite | 1,071 |
| Omics Experiment | 726 |

**Figure 1:**
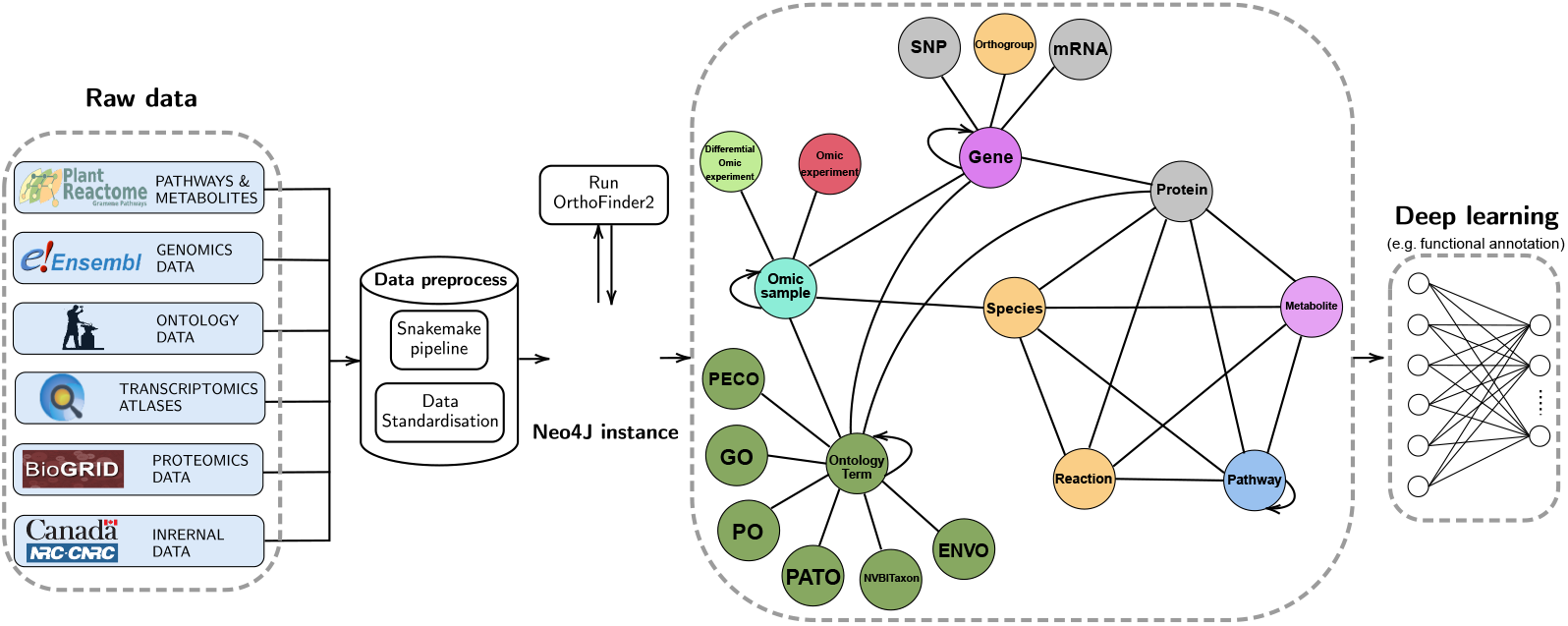
Complete pipeline including fetching of raw data from multiple data sources, incorporation of data into the DasDB, and parsing the graph data for training deep learning models for improving the functional annotation.

### 2.2 Predicting functional annotations from graph database

#### Knowledge Graph Reasoning

Our multi-omic graph database is considered as a knowledge graph *𝒢* = (*𝒱, ℰ, 𝒞, ℛ*), where *𝒱, ℰ* represent sets of entities and edges respectively, while 𝒞 and *ℛ* indicate sets of all node and edge types. Each node *u* ∈ 𝒱 corresponds to a label from set *𝒞*, such that *u* ∈ 𝒱_*c*_ where *c* ∈ 𝒞. On the other hand, each edge *e ∈ 𝒱* _*r*_ *∈ 𝒱*such that *r ∈ ℛ*. The knowledge graph *𝒢* is constructed from a set of facts, in which each fact is a triplet (*x, r, y*) *∈𝒱 ×ℛ × 𝒱* that indicates the relations *r* from node *x* to node *y*. Given the queries (*u, q*, ?), knowledge graph reasoning aims to find a subset of nodes *𝒱*_?_ *⊆ 𝒱*, such that *∀v ∈ 𝒱*_?_, the fact triplet (*u, q, v*) is true. In this work, we consider our multi-omic graph data as a knowledge graph and aim to predict related functional annotations represented in the knowledge graph given a gene. In particular, we focus on answering the query (*u, q, v*), such that *u ∈ 𝒱*_gene_, *v ∈ 𝒱*_functional-annotation_ and *q* is the relation between *u* and *v*.

#### Model selection

In this study, we aim to determine whether the graph topology of a multi-omic graph can contribute to predicting related functional annotations of given genes. We select a GNN model from the literature according to the following criteria: i) state-of-the-art GNNs that explore the topological aspect of the graph to perform knowledge graph reasoning; ii) high scalability to adapt to massive multi-omic graph data; iii) explainability to support prediction interpretation and clarification; and iv) ability to perform in the inductive setting (i.e reasoning on new species). There has been a surge in the number of studies in the past decade focusing on knowledge graph completion. The majority of proposed methods can be classified into three main paradigms (Zhu et al., 2021): path-based methods (Lao and Cohen, 2010; Gardner and Mitchell, 2015), embedding methods (Perozzi et al., 2014; Tang et al., 2015; Bordes et al., 2013a; Yang et al., 2015; Zhou et al., 2021) and GNNs (Kipf and Welling, 2016; Schlichtkrull et al., 2018; Davidson et al., 2018; Vashishth et al., 2020; Teru et al., 2020; Zhang and Chen, 2018). While path-based methods can be interpreted based on paths found by an existing model, this method requires the user to carefully hand-craft the reward function (Lin et al., 2018) or search strategy (Shen et al., 2018). On the other hand, embedding methods only encode node pairs, preventing them from reasoning on new species with a new set of nodes. Existing GNN methods are proven to be powerful when applied in an inductive setting – however, many methods are not scalable to large graphs and are limited in interpretability. Combining interpretability from path-based methods and the ability to perform reasoning on GNNs, A*Net is a scalable, path-based GNN that achieves state-of-the-art performance on knowledge graph reasoning Zhu et al.. Therefore, we use A*Net as an example model in this study.

#### A*Net

Knowledge graph completion aims to predict query relation *q* between source entity *u* and entity *v*. From the representation learning perspective, this requires learning a representation of a pair of entities *h*_*q*_(*u, v*) (Zhu et al., 2021). Inspired by traditional methods in link prediction, such as Katz index, Personalized PageRank, graph distance, widest path and most reliable path, A*Net formulates the pair representation as a *generalized sum* of all representations of paths between two entities *u* and *v* with a commutative summation operation ⊕, where each path representation *h*_*q*_(*P*) is defined as *generalized product* of all edge representations in the path with multiplication operation ⊗ (Zhu et al., 2021).

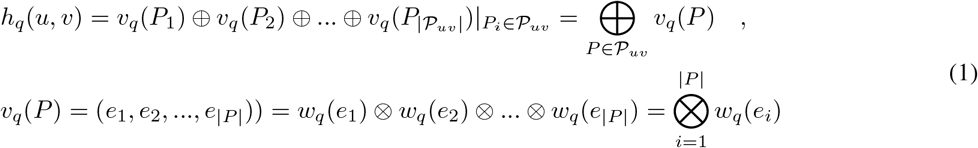

where *𝒫*_*uv*_ indicates a set of paths from *u* to *v* and *w*_*q*_(*e*_*i*_) is the presentations of edge *e*_*i*_. ⊕ and ⊗ are operations that generalize traditional path formulation algorithms ⊕ = +,⊗ = *×* for Katz index and Personalized PageRank; ⊕ = *min, ⊗* = + for Graph Distance; ⊕ = *max, ⊗*= *min* for widest path; ⊕= *max*, ⊗ = *×* for most reliable path (Zhu et al., 2021). To reduce the time complexity of enumerating all possible paths, A*Net adopts an approach that iteratively propagates the representations of t− 1 hops to nodes with higher priority according to a heuristic function to compute the representations of *t* hops (Hart et al., 1968). For a detailed description of the model’s architecture and implementation, we refer to the A*Net paper (Zhu et al., 2022).

#### Implementation

We use the implementation of A*Net provided by its authors (Zhu et al., 2022). All hyperparameters remain as described by Zhu et al. (2022). We train A*Net on 8 NVIDIA A100 GPUs (48GB) for 20 epochs and select the best model based on validation performance.

#### Functional annotation evaluation

We evaluate performance using a standard metric, mean reciprocal rank (MRR). MRR is a ranking quality metric that considers the position of the first relevant prediction in a ranked list. Given the *i*^*th*^ query (*u, q*, ?)^*i*^, the prediction provided by the model is a set of nodes 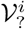, sorted in descending order according to the weight of each node 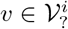. The MRR is computed as follows:

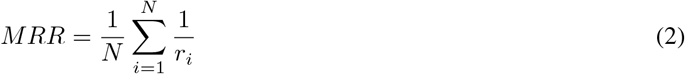

Where *N* is the total number of queries, *r*_*i*_ is the relevant position of the first valid related functional annotation of a gene of query *i*^*th*^. MRR values range from 0 to 1, where 1 indicates that the first relevant functional annotation is always at the top, and a higher MRR means better model performance. In addition to MRR, we evaluate the quality of models’ predictions by computing the ratio of related functional annotations of a gene among the top *N* predictions, *qual*@*N*. Higher *qual*@*N* indicates a higher probability of the model predicting correct related functional annotations given a gene.

#### Prioritize functional annotation scores

As the A*net generates a score that is different for each predicted edge in the function of the source node, additional heuristic manipulation is needed to interpret the importance or relevance of each target node, in our case, a predicted functional annotation of the source gene node. A*Net assigns a priority score to each node in the knowledge graph, reflecting its importance for answering a specific query. Given its relationship with the source gene, the neural priority function computes this score for each functional annotation. The GO terms with higher A*Net scores will be selected, indicating greater relevance to the query gene. A higher score from the neural priority function, *s*(*t*)*uq*(*x*), suggests that the predicted functional annotation is more likely to be part of an important path connecting the gene to relevant biological functions. The scores are evaluated in the context of the overall score range (e.g. Min: -0.777, Max: 0.207). A score close to the maximum (e.g. 0.207) would suggest a strong association. In contrast, a score closer to the minimum (e.g., -0.777) would indicate a weak or negative association between the source gene and the functional annotations. A heuristic threshold score was set, above which, a predicted functional annotation was considered significantly relevant. The functional annotation was selected as significant if the score is above 0.0 or in the first 50% of the score range.

### 2.3 Validation of the predicted functional annotation using gene expression

#### Collecting single-cell expression from *Arabidopsis thaliana* for validations

To validate the A*Net model prediction, we used gene expression data as a signature of functional annotation. The single-cell data were collected from the scPlantDB resource (He et al., 2023). Four published datasets that were formatted in scPlantDB were selected for their complementarity to represent a full view of Arabidopsis tissue types (Neumann et al., 2022; Kim et al., 2021; Liu et al., 2020; Shahan et al., 2022). The data were collected as R Seurat objects. The data were combined, normalized, and scaled.

#### Comparison of expression patterns

For each Gene Ontology (GO) or each Plant Ontology (PO) found in *Arabidopsis thaliana*, the expressions of the genes known to be part of this annotation were averaged. The same expression averages were made for the predicted genes for the same annotation. The Pearson Correlation Coefficient (PCC) was established between known and predicted gene expression averages. The PCC was done for all annotations in *Arabidopsis thaliana*, providing a comprehensive overview. The PCCs distribution from strong positive to negative correlation was used to control the A*Net overall prediction in *Arabidopsis thaliana*.

### 2.4 Discovery of pea nodule regulators of Nitrogen metabolism

#### Nodule nitrogen related genes extraction

The DasDB was used to extract the regulators of the pea nodule and the key nitrogen metabolism regulators. The query in Table 2 was performed to extract putative pea orthologs of the *Arabidopsis thaliana* genes involved in nitrogen metabolism. Furthermore, these genes have to be highly expressed in the nodules, the root organ of nitrogen uptake from the Rhizobium, symbiotic bacteroids transforming atmospheric nitrogen into an available nitrogen source for the plant.

**Table 2:**
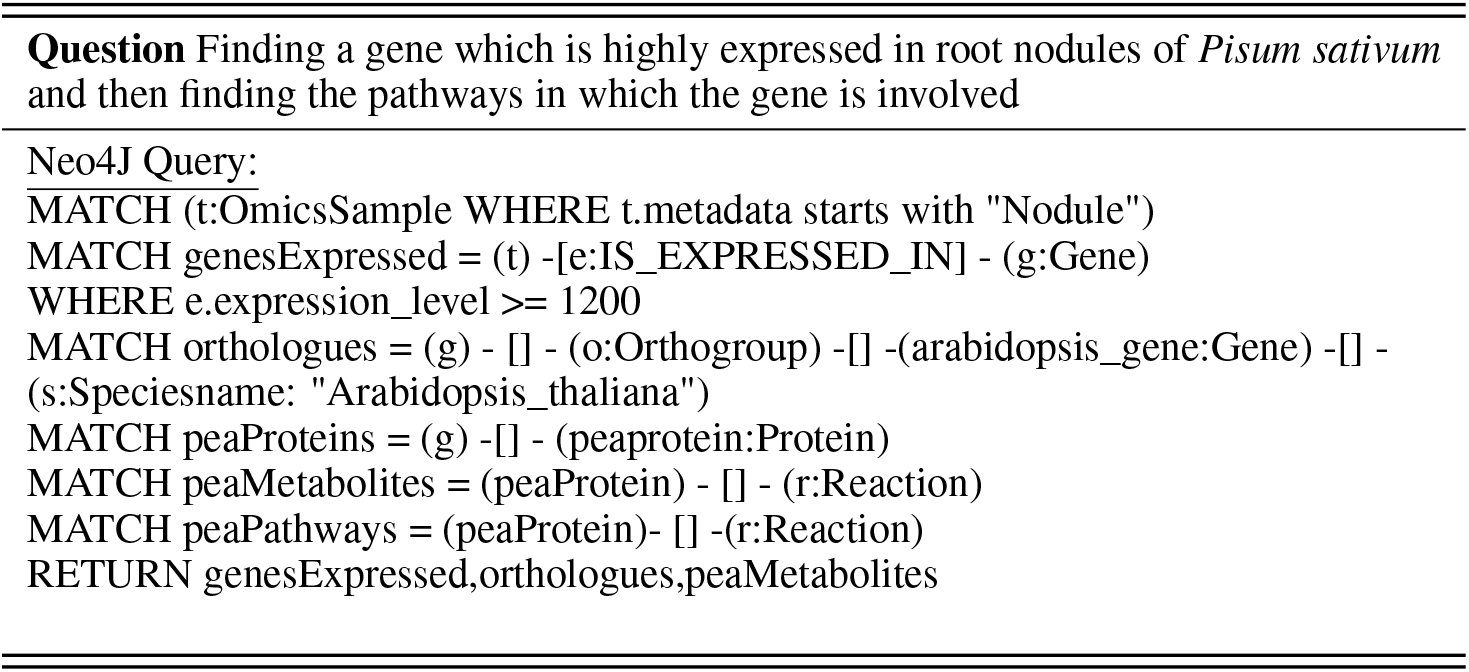
Retrieve meaningful information from multi-omic database.

#### Regulators discovery

The above query on nodule nitrogen enables the extraction of a multi-node type network. To find the putative regulators of this network in pea, the PageRank from the NetworkX Python library was performed. Only the pea genes with the highest PageRanks were retained for further analysis on the regulators.

#### Comparison of known vs predicted annotations

The known annotations for pea genes were extracted from the DasDB using the following query:

~~~
MATCH (s:Species\{name:”Pisum\_sativum”\})-[]-(g:Gene)-[]-(a:n4sch\_\_Class)
RETURN g.name, a.n4sch\_\_name}.
~~~

Then, the table of new predictions obtained for the nitrogen sub-network was merged with the known annotations, and gene definitions were created using a Python custom script.

## 3 Results

### 3.1 DasDB multi-omic database

The DasDB contains 17 plant species: *Arabidopsis thaliana*, Oil rapeseed (*Brassica napus*), field mustard (*Brassica rapa*), soybean (*Glycine max*), barrel clover (*Medicago truncatula*), pea (*Pisum sativum*), chickpea (*Cicer arietinum*), lentil (*Lens culinaris*), common bean (*Phaseolus vulgaris*), common grape vine (*Vitis vinifera*), faba beans (*Vicia faba*), mung bean (*Vigna radiata*), camelina (*Camelina sativa*), rice (*Oryza sativa*), maize (*Zea mays)*, wheat (*Triticum aestivum*), Barley (*Hordeum vulgare*) and the great millet (*Sorghum bicolor*). The 2,620,206 functional annotations extracted from the OBO Foundry were represented by Gene Ontology (GO), Plant Ontology (PO), and Plant Experimental Conditions Ontology (PECO).

### 3.2 Evaluation Methodology

#### Datasets and Evaluation

We evaluate the multi-omic graph database, the DasDB, produced by the pipeline discussed in Section 2.1, on various functional annotation tasks. For the first experiment, we obtain from the graph database a subgraph capturing the information about the genome, pathway, proteome, orthogroup and phenotypes and their relations in a well-studied plant species, *Arabidopsis thaliana*. Then, we performed functional annotation prediction by reasoning the sub-knowledge graph by training the A*Net. In the first experiment, we split all relations between genes and phenotypes into train, validation, and test splits with a ratio of 90*/*5*/*5.

For the second experiment, we evaluate transferability in predicting phenotype from a well-studied species to a novel species, which is practical in many real-world applications. To facilitate the second experiment, we use a subset of species, including *Arabidopsis thaliana, Zea mays, Oryza sativa* and *Sorghum bicolor*, and obtain a single unified graph containing genome, pathway, proteome, orthogroup and functional annotations and their relations in those species. In contrast to the first experiment, we conducted each training, validation and testing on different species in the second experiment (e.g. Training on *Arabidopsis thaliana, Zea mays*, Testing on *Oryza sativa* and Validation on *Sorghum bicolor*).

In both experiments, most links between functional annotations and genes are excluded from the graph, and the model is asked to predict those links to avoid information leakage. This leakage occurs when the model can access target relationships during training, allowing it to memorize them instead of learning to predict them, leading to overly optimistic performance. However, a few links are kept to avoid disconnecting the graph. In addition, we augment each triplet (*u, q, v*) with a flipped triplet (*v, q*^*−*^1, *u*) in the preprocessing step as previously described (Zhu et al., 2021, 2022). For evaluation, we use the standard filtered ranking protocol (Bordes et al., 2013a) for the knowledge graph, where each triplet (*u, q, v*) is ranked against all negative triplets (*u, q, v*^*′*^) or (*u*^*′*^, *q, v*) that are not present in the knowledge graph. We measure the performance with mean reciprocal rank (MRR) and the proportion of ground true predictions among the top *k* predictions given from the models, termed HITS at K (*hits*@*k*). We report the average and standard deviation performance on three different random splits of the data.

#### Baselines

As a naive baseline, we used a static list of related phenotypes for all genes according to the commonality among genes in the training set (Common rank). For example, the first entry in the list is the gene related to the largest number of functional annotations in the training set. As additional baselines, we also test embedding methods TransE (Bordes et al., 2013b), DistMult (Yang et al., 2014), ComplEx (Trouillon et al., 2016), and RotatE (Sun et al., 2019), which learn a distributed representation for each node and edge by preserving the edge structure of the graph. While embedding methods can achieve promising results on knowledge graph completion, they do not explicitly encode local subgraphs between node pairs, leading to limited performance on new species.

### 3.3 Predicting Functional Annotations

#### Predicting gene annotations on an *Arabidopsis thaliana* multi-omic graph

We first evaluate the ability of the model to predict functional annotations on a well-studied species, *Arabidopsis thaliana*. (Table 3). A*Net outperforms the static baseline and the embedding-based methods in MRR, *qual*@1, and *qual*@3. Because the primary difference between A*Net and the baseline methods is the former’s ability to reason using graph topology, we attribute the success of GNN-based models to their ability to learn the structure of comprehensive interaction networks of multi-omic data.

**Table 3:** Train and test on *Arabidopsis thaliana*.

| Model | MRR | hits@1 | hits@3 | hits@10 | hits@20 | hits@50 |
| --- | --- | --- | --- | --- | --- | --- |
| A*Net | **0.723 ± 0.097** | **0.674 ± 0.097** | **0.616 ± 0.144** | 0.567 ± 0.163 | 0.462 ± 0.141 | 0.273 ± 0.083 |
| Common rank | 0.671 ± 0.195 | 0.520 ± 0.294 | 0.567 ± 0.237 | **0.593 ± 0.201** | **0.572 ± 0.173** | 0.334 ± 0.081 |
| TransE | 0.420 ± 0.120 | 0.278 ± 0.115 | 0.296 ± 0.136 | 0.323 ± 0.151 | 0.372 ± 0.15 | 0.241 ± 0.092 |
| RotatE | 0.703 ± 0.030 | 0.580 ± 0.036 | 0.532 ± 0.036 | 0.447 ± 0.019 | 0.408 ± 0.010 | 0.271 ± 0.010 |
| DistMult | 0.636 ± 0.095 | 0.442 ± 0.166 | 0.498 ± 0.051 | 0.485 ± 0.023 | 0.481 ± 0.049 | 0.319 ± 0.050 |
| ComplEx | 0.701 ± 0.158 | 0.558 ± 0.233 | 0.515 ± 0.121 | 0.576 ± 0.148 | 0.570 ± 0.136 | **0.337 ± 0.068** |

To evaluate whether the information from multiple omics contributes to the model’s performance, we conduct an ablation study where we train the models on the datasets where all information has been removed except for genes and phenotypes. Training A*Net on the dataset containing only genes and phenotypes yielded universally worse performance (Table 4). The gaps between the performance of the models trained in two settings are significant when it comes to *hits*@3, *hits*@10, *hits*@20 and *hits*@50, which highlights the advantages of multi-omic data in biological tasks, especially for annotating genes.

**Table 4:** Ablation study using only gene and phenotype information (A*Net on *Arabidopsis thaliana*)

| Setting | MRR | hits@1 | hits@3 | hits@10 | hits@20 | hits@50 |
| --- | --- | --- | --- | --- | --- | --- |
| Completed graph | **0.723 ± 0.097** | **0.674 ± 0.097** | **0.616 ± 0.144** | **0.567 ± 0.163** | **0.462 ± 0.141** | **0.273 ± 0.083** |
| Only genes & phenotypes | 0.717 ± 0.26 | 0.635 ± 0.349 | 0.311 ± 0.162 | 0.228 ± 0.087 | 0.185 ± 0.076 | 0.134 ± 0.035 |

#### Transferability from *Arabidopsis thaliana* to *Sorghum bicolor*

We explored the transferability of GNN-based models and multi-omic data across species by training and testing on different species. An extensive number of works have been studying model species such as *Arabidopsis thaliana*, resulting in a large quantity of high-quality data on these species, while information on other species remains sparse. Therefore, the transference of functional information from well-resourced to under-resourced species is a valuable application for predictive modelling. We evaluated the transferability of our approach by training the models on two species, including *Arabidopsis thaliana* and *Zea mays*, and testing on a third species, *Sorghum bicolor* (Table 5). Across all metrics, the proposed approach was the only one among the tested ones capable of transferring information across species. The performance of all baselines significantly decreased in the transferability setting compared to training and testing on the same species.

**Table 5:** Transferability test of the models. The models are trained on two species and then tested and validated on the other two species.

| Setting | MRR | hits@1 | hits@3 | hits@10 | hits@20 | hits@50 |
| --- | --- | --- | --- | --- | --- | --- |
| A*Net | **0.606 ± 0.109** | **0.509 ± 0.155** | **0.446 ± 0.122** | **0.300 ± 0.063** | **0.196 ± 0.044** | **0.102 ± 0.023** |
| Common rank | 0.096 ± 0.011 | 0.000 ± 0.000 | 0.049 ± 0.000 | 0.024 ± 0.000 | 0.012 ± 0.000 | 0.033 ± 0.000 |
| TransE | 0.067 ± 0.064 | 0.024 ± 0.050 | 0.017 ± 0.026 | 0.015 ± 0.012 | 0.013 ± 0.007 | 0.013 ± 0.005 |
| RotatE | 0.089 ± 0.022 | 0.023 ± 0.009 | 0.026 ± 0.014 | 0.032 ± 0.014 | 0.036 ± 0.007 | 0.037 ± 0.007 |
| DistMult | 0.088 ± 0.095 | 0.030 ± 0.047 | 0.039 ± 0.052 | 0.040 ± 0.048 | 0.033 ± 0.036 | 0.021 ± 0.021 |
| ComplEx | $1.7 \times 10^{-6} \pm 2.2 \times 10^{-7}$ | 0.000 ± 0.000 | 0.000 ± 0.000 | 0.000 ± 0.000 | 0.000 ± 0.000 | 0.000 ± 0.000 |

#### Prediction interpretation

One advantage of the studied model is that we can interpret its predictions through paths in the knowledge graph. This may provide insight for biologists to interpret and clarify the model predictions. Intuitively, the interpretations should contain paths that contribute most to the prediction *p*(*u, q, v*). The example of the model’s explanation of the prediction is shown in Table 6.

**Table 6:**
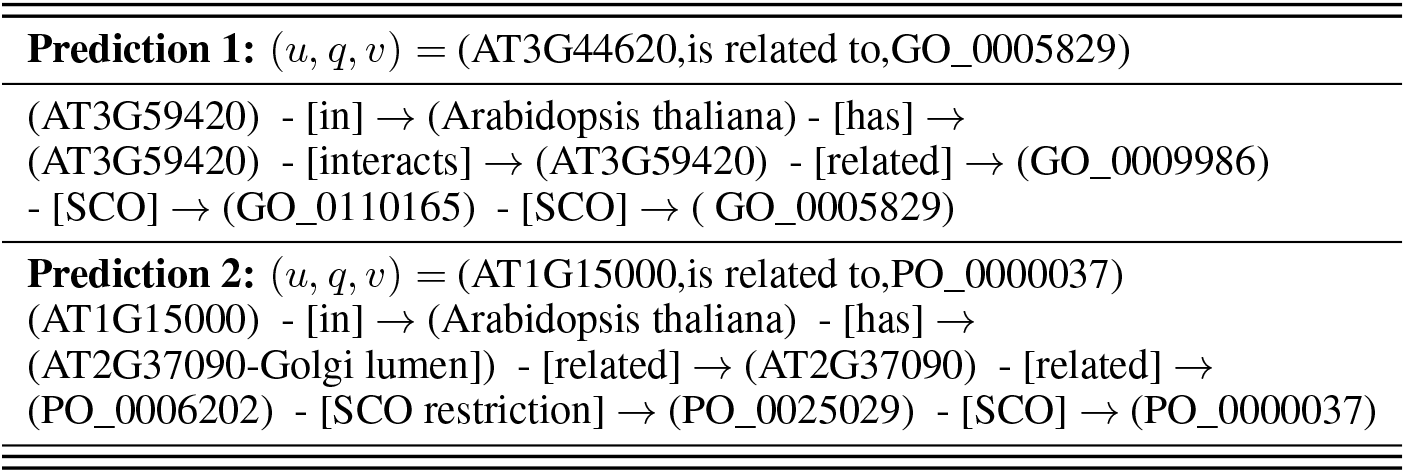
Model’s prediction interpretation.

### 3.4 Model validation using gene expression patterns (Comparing known vs predicted functional annotations.)

A large portion of single-cell data published in *Arabidopsis thaliana* was normalized and grouped in an integrated dataset (He et al., 2023). The single-cell data grouped in pseudo-replicates were used to study the transcript expression mean patterns for genes falling under each functional annotation category using GO (Gene Ontology) and PO (Plant Ontology) annotations. Average gene expression patterns were calculated for each GO annotation for the known and the predicted GO annotations from the A*Net results without empirical filtering of the GNN score.

Pearson correlation coefficients were then calculated between the mean expression values of new predictions and known annotations for each category. These coefficients indicated the strength and direction of the linear relationship between the two datasets for each category.

The correlation coefficients were categorized into different ranges (e.g., Strong Negative to Very Strong Positive correlations) to understand better the relationship between the expression patterns for genes known and predicted to be part of a functional annotation, as shown in Figure 2. The frequency of each correlation range was summarized and visualized in a bar plot (Figure 3). Showing the distribution of Pearson correlation coefficients across different ranges provided insights into the agreement between new predictions and known annotations. Of 6674 functional annotations, 38 had no gene expression pattern attached to them (Supplemental Table S1). No annotation correlated negatively with the expression of its known vs predicted genes. 53% (3575/6636) of the correlations between known and predicted were Strong Positive or Very Strong Positive correlations (between 0.7 and 1 Pearson Correlation). Adding the Moderate Positive brought the relationship to 70%. This result tends to confirm experimentally the accuracy of the A*Net model in predicting accurate functional annotations using gene expression patterns as a proxy of functional annotation signatures.

**Figure 2:**
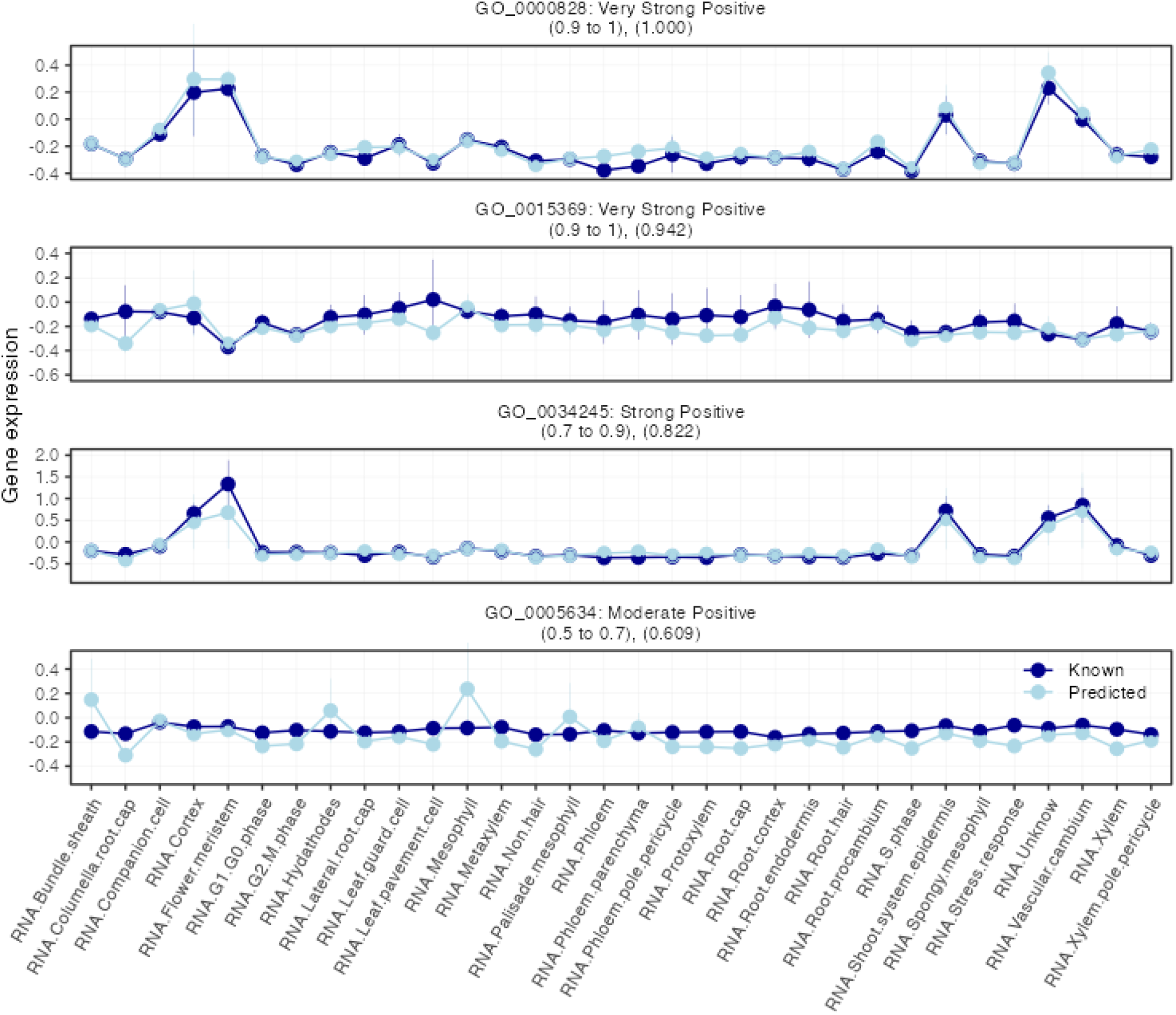
Example of gene expression profiles for predicted versus known annotations using the *Arabidopsis thaliana* single-cell data

**Figure 3:**
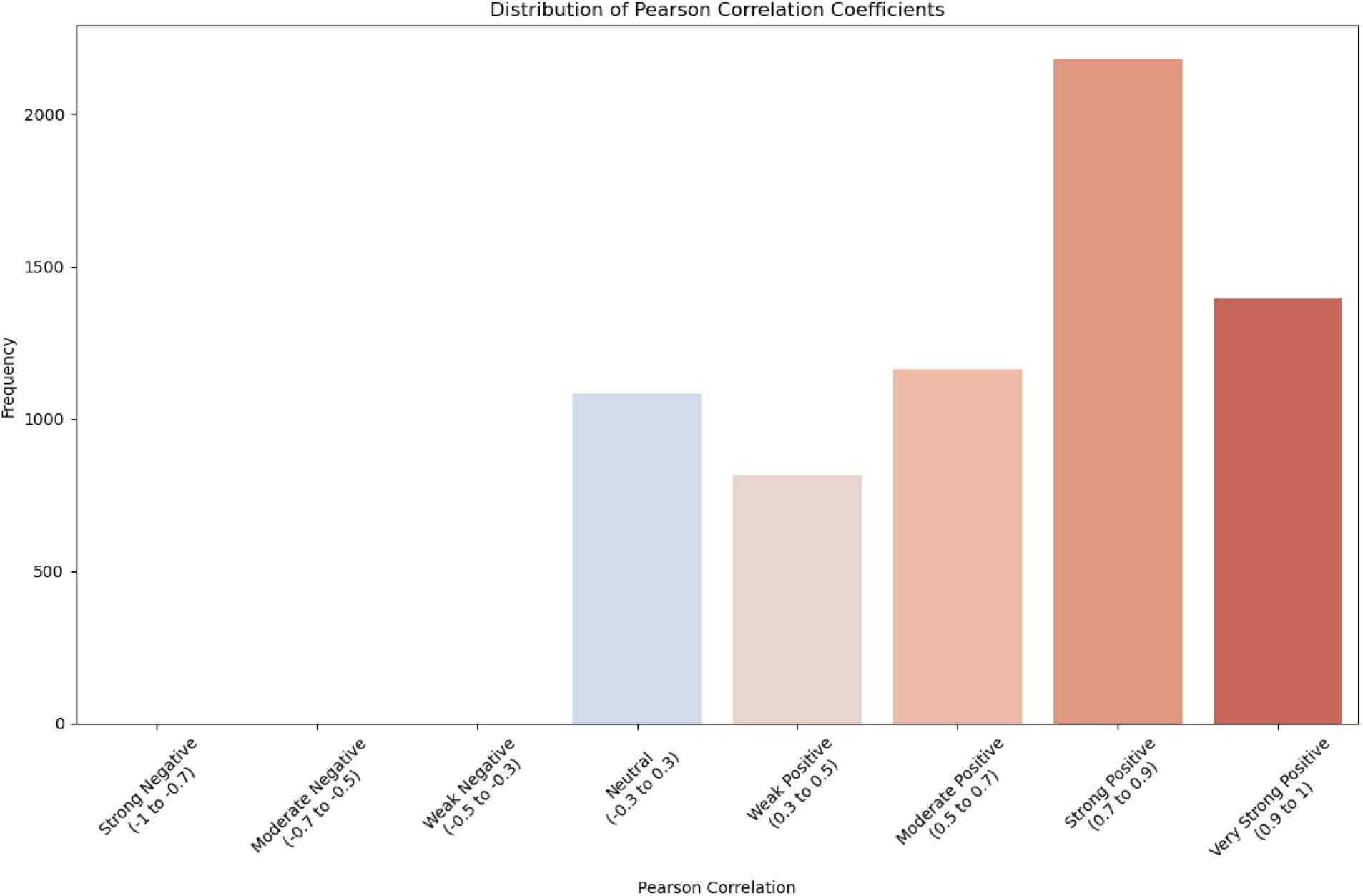
Comparison of gene expression profiles for predicted versus known annotations

### 3.5 Pea genome functional annotation

There are 3,361 (9,077 - 5,716) genes annotated by the A*Net model functional annotation prediction, having previously no annotation that could be transferred from the orthology from Arabidopsis to Pea Table 7. These genes are not annotated in the Pea genome or in the Arabidopsis genome. The A*Net model functional annotation prediction is a new prediction that is not based on the orthology transfer. These annotations came from other species or the short links with other annotations.

**Table 7:** Annotation in Pea genome generated from the A*Net model functional annotation prediction in *Pisum sativum*.

| Category | Count |
| --- | --- |
| No Annotation before | 9,077 |
| No Annotation after | 5,716 |
| Total Pea Genes | 31,340 |

The A*Net model functional annotation prediction has predicted additional annotations not based on orthology transfer for 24,919 genes Table 8. Only 369 of these genes received annotations already present in the orthology transfer Table 8. These latter annotations can represent the limited leakage of the model described in the method section. 398,783 new annotations were assigned to these 25k genes Table 8.

**Table 8:**
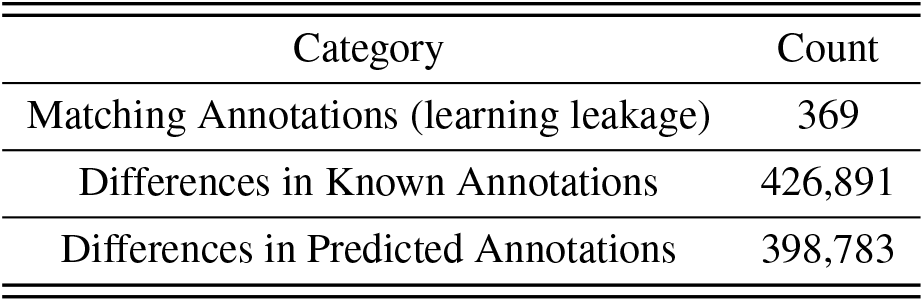
Annotation in Pea genome generated from the A*Net model functional annotation prediction in *Pisum sativum*.

### 3.6 Biological cases: Analysis of the Sub-network of Nodule Genes Linked to Low Nitrogen Conditions in Pea

Using the DasDB knowledge graph, the nitrogen-related sub-network expressed in pea root nodules was extracted. A sample of this sub-network was represented to illustrate the organization of the extracted data (Figure 4). This sample showed four pea genes: Psat2g056920, Psat5g153000, Psat7g019040, Psat6g079280. They are the putative orthologs of nitrogen-related Arabidopsis genes. The three transcriptomic samples extracted from the query were aeroponic and hydroponic in low nitrate treatments (SRR1693197, SRR1691099, and SRR1692096). These omics samples were part of the experiment PRJNA267198 (https://www.ncbi.nlm.nih.gov/bioproject/267198) in the Cameor pea lines (Kreplak and Burstin, 2014).

**Figure 4:**
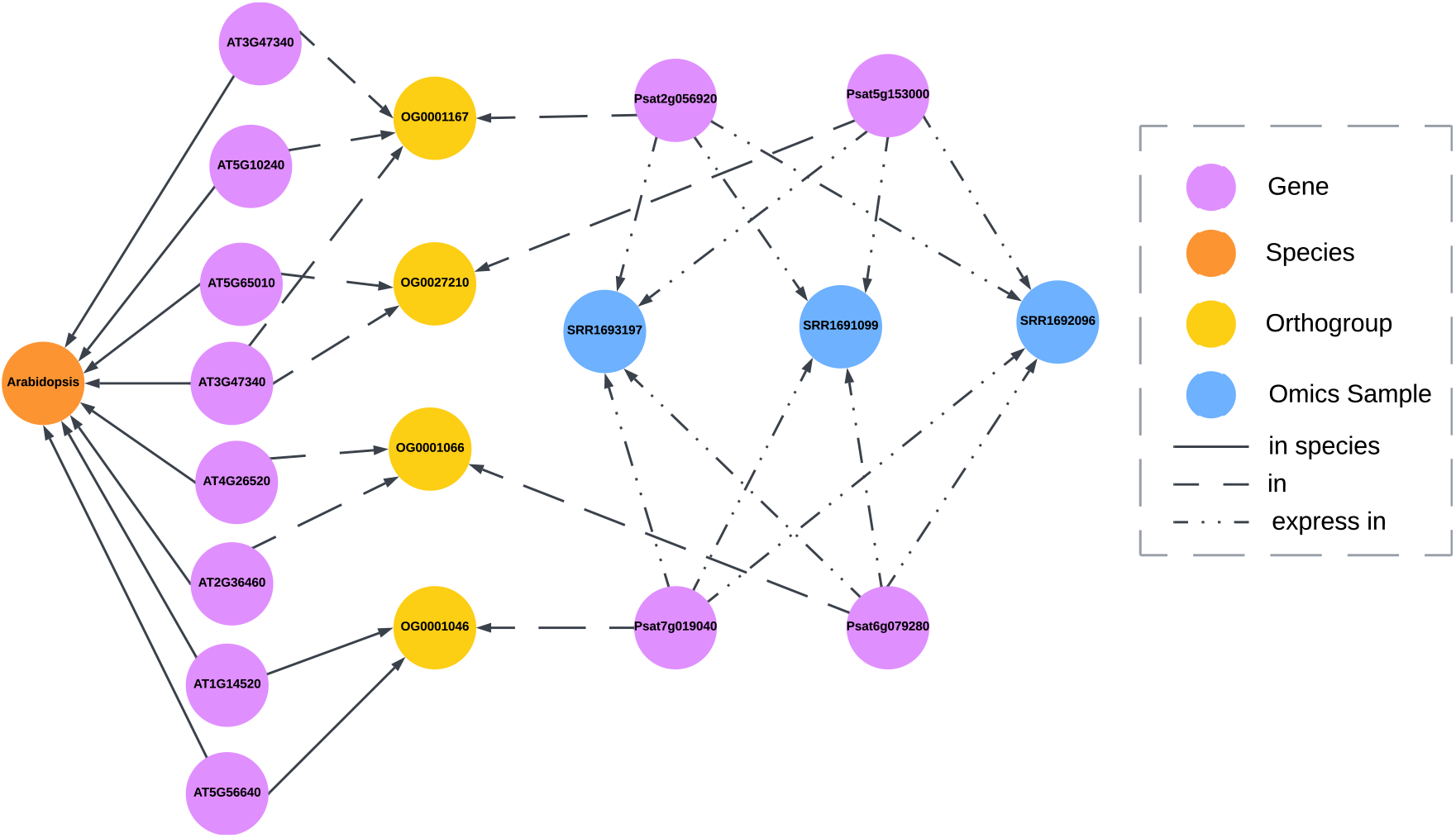
Results obtained from query described in Table 2 to multi-omic graph database

#### 3.6.1 Comparison with Existing Functional Annotations *(Overlap with known databases, GO term enrichment.)*

Our results were compared with existing functional databases such as Gene Ontology (GO) and Plant Ontology (PO) to ensure the accuracy and biological relevance of the predicted functional annotations. This comprehensive analysis, detailed in Supplemental Table S2, revealed that eight genes out of the 23 in the sub-network with high PageRank had a redundant annotation with the known ones. These genes were involved in core metabolic process pathways. The high PageRank scores of genes showed strong enrichment in known functional categories essential for nitrogen uptake in pea nodules, reinforcing the importance of the selected biological networks. We could identify that the genes identified followed the N/C nutrient process in pea root nodules as identified in Figure 5.

**Figure 5:**
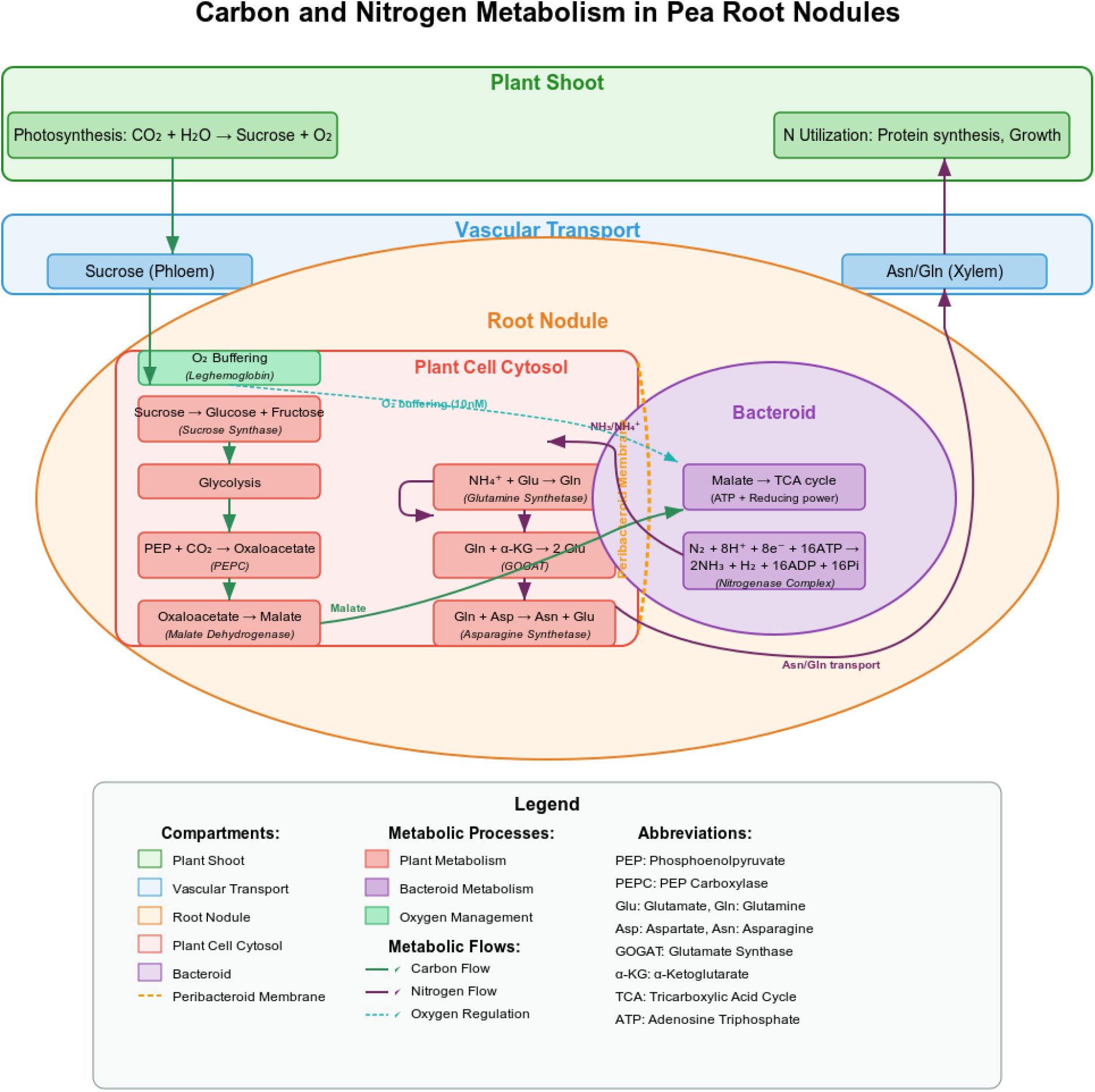
Carbon and nitrogen metabolism in pea root nodules

#### 3.6.2 Key genes identification

To further illustrate the impact of our predicted functional annotations, we examined case studies of specific genes that were highly ranked in our network analysis and how the A*Net model improved their functional characterizations. These case studies highlight the potential biological roles of these genes and provide a framework for future experimental validation. The PageRank algorithm was applied to prioritize genes based on their importance within the network. Many of the high-ranking genes were previously annotated in biological databases, validating the effectiveness of our predictive model. Additionally, we identified novel high-PageRank genes without prior annotations, highlighting potential targets for further functional characterization.

##### High-PageRank Gene Candidates in Nitrogen Metabolism

Several top-ranked genes identified using PageRank exhibited strong connectivity with known nitrogen metabolism genes. One candidate gene (Psat2g056920, glutamine-dependent asparagine synthase 1) correlated highly with nodulation-related genes and was differentially expressed under low nitrogen conditions. This suggests a potential regulatory role in nitrogen uptake or assimilation, warranting further investigation through experimental validation. Some of these genes, such as Psat0s690g0040 and Psat5g120440, were involved in nitrogen uptake. The homologues of glutamate-ammonia ligase were part of forming L-glutamate, the pathway assimilating nitrogen in amino acids, after the absorption of ammonia (*NH*_3_) (R-R-PSA-1121516). *NH*_3_ is a product released by the rhizobium from atmospheric nitrogen transformation. Other genes involved in nitrogen assimilation and incorporation in amino acid metabolism, such as another glutamine-dependent asparagine synthase 1 (Psat5g153000), amino acid synthesis genes, were also identified.

##### High-PageRank Gene Candidates in Carbon Metabolism

Cytosolic glycolysis is one of the main pathways central to the highly expressed genes in the nodules (R-PSA-1119570). A fructose-bisphosphate aldolase (Psat6g079280) transforms the fructose bisphosphate into GAP3P (D-glycerate 3-phosphate) and DHAP (dihydroxyacetone phosphate). The next step in the pathway is the catalysis of GAP3P by glyceraldehyde-3-phosphate dehydrogenase. Two homologues of glyceraldehyde-3-phosphate dehydrogenase, Psat5g094000 and Psat2g071480, were also part of the most central genes of the nitrogen-related nodule sub-network. The production of sucrose (R-ALL-29532) was hypothesized in the nodule due to the high expression of sucrose synthase (Psat4g019440), another gene of carbon metabolism central in the nitrogen-related nodule sub-network. In nodules, sucrose was shown to be crucial in nitrogen fixation and respiration processes of the bacteroid through the leghemoglobin protein (Gordon et al., 1999). Downstream to the sucrose metabolism, a homologue of inositol oxygenase (Psat7g019040) was highly ranked. The inositol oxygenase enzyme transforms inositol into D-glucuronate, part of the UDP-D-glucuronate biosynthesis pathway (Reactome: R-PSA-1119431), as shown in Figure 5.

##### High-PageRank Gene Candidates in Symbiote Respiration

A homologue of carbonic anhydrase (Psat0s2720g0080), part of the cyanate degradation pathway (R-PSA-1119586), was within the nitrogen-related nodule sub-network. Carbonic anhydrases catalyze the reaction of *CO*_2_ with water to produce bicarbonate ions (*HCO*_3_^*−*^), which can be readily utilized by the bacteria inside the nodules (Kavroulakis et al., 2000). Carbonic anhydrases were specifically expressed in the nodule tissues, particularly in the cells surrounding the bacteria (Gálvez et al., 2000). By providing a carbon source in the form of bicarbonate, carbonic anhydrase plays a vital role in supporting the energy requirements of nitrogen fixation by the bacteria (Wang et al., 2021a).

##### Novel Genes in Nodule Development

Several predicted genes were highly connected to known nodulation regulators but lacked prior GO annotations linked to nitrogen or nodule processes. One aquaporin (Psat7g067400) was one of the most critical genes expressed in nodules under low nitrogen conditions (R-PSA-9618197-8). It displayed strong enrichment in pea root nodules and was co-expressed with genes involved in early nodule formation. This gene’s high PageRank degree of centrality suggests it may play a crucial role in the signaling cascade leading to nodule organogenesis. This aquaporin is a homolog of NIP1;1 and NIP3;1, NOD-like transporters known to be linked to detoxification in Arabidopsis (Wang et al., 2017, 2020). Interestingly, glyceraldehyde-3-phosphate dehydrogenase (Psat5g094000) has a predicted annotation linked to aerobic respiration (GO_0019411). The respiration process is crucial for the bacteroids’ survival and is regulated by the carbon metabolism in nodules. Our functional annotation has putatively generated a link between carbon metabolism and the host provision of a low aerobic environment for the bacteroid through the leghemoglobin protein (Gordon et al., 1999), as shown in Figure 5.

##### Predicted Bacteroid-containing Symbiosome

Among the most central genes of the nitrogen-related nodule sub-network, four genes were linked to the annotation GO_0043660, which describes a bacteroid-containing symbiosome. These genes included those known as carbon metabolism components (Psat4g019440, sucrose synthase) and nitrogen metabolism (Psat2g056920, asparagine synthase), a heme-like binding protein potentially important for the respiration of the symbiotic bacteroid (Psat2g053680), and an unknown protein linked to cell membrane trafficking according to the predicted annotation (Supplemental Table S2).

## 4 Discussions

A knowledge graph, the DasDB, was developed for crop data. This tool presents a pragmatic approach to connect several data sources, from transcriptomics to functional annotation. The DasDB is not as comprehensive as KNetMiner (Hassani-Pak et al., 2021) or as specialized as the Ortho_KB (Imbert et al., 2023). However, the DasDB describes a framework to generate such knowledge graph methodology, which can be implemented quickly to answer specific questions and formulate hypotheses with limited computational resources. The graph database framework, DasDB, enables the easy and rapid expansion of the knowledge graph to other datasets and ensures the automatic integration of metadata in the node or relationship properties. Moreover, DasDB provides researchers with the assurance that their data will adhere to the FAIR principles, as the storage and properties of the nodes and relationships are automatically formatted accordingly.

The A*Net, a graph neural network, was implemented on top of the knowledge graph to understand better the network extracted from a query and increase the discovery power. The prediction produced by A*Net on Arabidopsis on functional annotation, was significantly higher than that of state-of-the-art embedding graph neural networks (Table 3). In addition, ablation studies highlighted the importance of the inclusion of multi-omic data for model performance (Table 4). However, the lack of initial data in a particular species made transferability less efficient for peas regarding the MRR score (Table 5 -0.606 pea transfer vs 0.723 intra-species - Arabidopsis). Another limitation of the Graph Neural Network is the tendency to generate biases to well-studied functional annotations (Gunning and Pavlidis, 2021). Our functional annotation model is a generalization on other attempts aiming to find new functional annotations for genes using a knowledge graph. Our functional annotation model is able to find the connection deep in the network not reachable by the former methods (Zhang et al., 2024).

The comparison with existing functional annotations confirms that our predictive approach successfully recapitulates known biological functions while identifying novel candidate genes for further investigation. Enriching predicted genes in biologically relevant pathways highlights the value of network-based annotation methods in advancing functional genomics research. We conducted a functional enrichment analysis using GO term enrichment tests to validate our predictions further. The study revealed significant enrichment in nitrogen metabolism, carbon metabolism, and nodulation categories, aligning with our focus on pea nodule expression and low nitrogen response. The presence of predicted genes in these enriched categories strengthens their potential biological relevance.

The network analysis of predicted functional genes reveals that genes with high connectivity and centrality measures are often involved in essential biological processes. These case studies illustrate the biological relevance of our network-based predictions by highlighting genes that show strong connectivity patterns but lack prior annotations. By prioritizing these genes for further experimental validation, our study provides a roadmap for uncovering novel gene functions in nitrogen metabolism, stress response, and nodule development. The integration of network analysis with biological case studies enhances the interpretability of our results and strengthens the application of graph-based methods in functional genomics research for complex systems.

Our examination of known and predicted functions has not only led to the discovery of several top-ranked genes associated with nitrogen metabolism, carbon and stress response pathways, but also validated our approach. These findings, which support their role in the pea nodule expression dataset, suggest that our approach effectively captures functionally essential genes. This may provide insights for future experimental validation.

### Novel Predictions and Potential Discoveries

In the pea example, beyond validating known functional annotations, our approach identified a subset of genes without prior annotations in GO or PO. These genes exhibited strong network connectivity and were frequently linked to biologically relevant pathways, suggesting potential novel functions. Further experimental validation of these genes could expand our understanding of their roles in plant development and stress adaptation. Consequently, new gene function hypotheses can be generated from the A*Net predicted annotations. For instance, two putative asparagine synthases 1 were present in the nodule nitrogen-related network. The predicted annotations showed that Psat5g153000 is related to root hair and trichoblast in the knowledge graph. Although expressed in nodules, Psat5g153000 could be associated with the early nodule and root nitrogen metabolism. The second putative asparagine synthase 1, Psat2g056920, was associated with the bacteroid-containing symbiosome annotation, showing a more specialized role in nodule nitrogen metabolism. For example, in soybean, the pea ortholog of asparagine synthase 1 was shown to be expressed in nodules with or without nitrate, an inhibitor of atmospheric nitrogen fixation (Antunes et al., 2008).

We have presented the pea nodule as an example of using DasDB and improving functional annotation through A*Net Graph neural network application. This well-characterized symbiotic process is key in crop rotation, improving soil health, and reducing the use of chemical fertilizers. The new targets identified in the study can be tested by selection in existing populations or creating them using SNP prediction algorithms (Frazer et al., 2021; Jumper et al., 2021) and gene editing techniques. This tool allows researchers to gain insights into a myriad of research questions, providing connections not available without the incorporation of multi-omic data resources. These insights can expedite research and aid in the development of crops with agronomically beneficial traits.

## Availability and implementation

The implementation of A*Net on the DasDB is available in the following GitHub repository: PlantBio KG reasoning

## Acknowledgments

We thank the Aquatic and Crop Resource Development for supporting T.G.B.N.’s and S.D.’s works. We thank Janice Schmidt for her guidance concerning the Ontology resources.

## References

D. Akhoundova and M. A. Rubin. Clinical application of advanced multi-omics tumor profiling: Shaping precision oncology of the future. Cancer Cell, 40(9):920–938, 2022.

F. Antunes, M. Aguilar, M. Pineda, and L. Sodek. Nitrogen stress and the expression of asparagine synthetase in roots and nodules of soybean (glycine max). Physiologia plantarum, 133(4):736–743, 2008.

A. Bordes, N. Usunier, A. García-Durán, J. Weston, and O. Yakhnenko. Translating embeddings for modeling multi-relational data. In C. J. C. Burges, L. Bottou, Z. Ghahramani, and K. Q. Weinberger, editors, Advances in Neural Information Processing Systems 26: 27th Annual Conference on Neural Information Processing Systems 2013. Proceedings of a meeting held December 5-8, 2013, Lake Tahoe, Nevada, United States, pages 2787–2795, 2013a. URL https://proceedings.neurips.cc/paper/2013/hash/1cecc7a77928ca8133fa24680a88d2f9-Abstract.html.

A. Bordes, N. Usunier, A. García-Durán, J. Weston, and O. Yakhnenko. Translating embeddings for modeling multi-relational data. In C. J. C. Burges, L. Bottou, Z. Ghahramani, and K. Q. Weinberger, editors, Advances in Neural Information Processing Systems 26: 27th Annual Conference on Neural Information Processing Systems 2013. Proceedings of a meeting held December 5-8, 2013, Lake Tahoe, Nevada, United States, pages 2787–2795, 2013b. URL https://proceedings.neurips.cc/paper/2013/hash/1cecc7a77928ca8133fa24680a88d2f9-Abstract.html.

Z. Cai, R. C. Poulos, J. Liu, and Q. Zhong. Machine learning for multi-omics data integration in cancer. Iscience, 25(2), 2022.

D. Chicco, F. Cumbo, and C. Angione. Ten quick tips for avoiding pitfalls in multi-omics data integration analyses. PLOS Computational Biology, 19(7):e1011224, 2023.

J. Confais and N. Francillonne. Linking heterogeneous data from model plant species in a graph database. In Séminaire résidentiel INRAE Semantic Linked Data édition 2023, 2023.

T. R. Davidson, L. Falorsi, N. D. Cao, T. Kipf, and J. M. Tomczak. Hyperspherical variational auto-encoders. In A. Globerson and R. Silva, editors, Proceedings of the Thirty-Fourth Conference on Uncertainty in Artificial Intelligence, UAI 2018, Monterey, California, USA, August 6-10, 2018, pages 856–865. AUAI Press, 2018. URL http://auai.org/uai2018/proceedings/papers/309.pdf.

D. M. Emms and S. Kelly. Orthofinder: phylogenetic orthology inference for comparative genomics. Genome biology, 20:1–14, 2019.

J. Frazer, P. Notin, M. Dias, A. Gomez, J. K. Min, K. Brock, Y. Gal, and D. S. Marks. Disease variant prediction with deep generative models of evolutionary data. Nature, 599(7883):91–95, 2021.

M. Gao, S. Liu, Y. Qi, X. Guo, and X. Shang. Gae-lga: integration of multi-omics data with graph autoencoders to identify lncrna–pcg associations. Briefings in Bioinformatics, 23(6):bbac452, 2022.

M. Gardner and T. M. Mitchell. Efficient and expressive knowledge base completion using subgraph feature extraction. In L. Màrquez, C. Callison-Burch, J. Su, D. Pighin, and Y. Marton, editors, Proceedings of the 2015 Conference on Empirical Methods in Natural Language Processing, EMNLP 2015, Lisbon, Portugal, September 17-21, 2015, pages 1488–1498. The Association for Computational Linguistics, 2015. doi: 10.18653/V1/D15-1173. URL https://doi.org/10.18653/v1/d15-1173.

J. Gillis and P. Pavlidis. The Impact of Multifunctional Genes on “Guilt by Association” Analysis. PLoS ONE, 6(2): e17258, 2011a. doi: 10.1371/journal.pone.0017258.

J. Gillis and P. Pavlidis. The role of indirect connections in gene networks in predicting function. Bioinformatics, 27 (13):1860–1866, 2011b.

P. Gong, L. Cheng, Z. Zhang, A. Meng, E. Li, J. Chen, and L. Zhang. Multi-omics integration method based on attention deep learning network for biomedical data classification. Computer Methods and Programs in Biomedicine, 231: 107377, 2023.

A. J. Gordon, F. R. Minchin, C. L. James, and O. Komina. Sucrose synthase in legume nodules is essential for nitrogen fixation. Plant physiology, 120(3):867–878, 1999.

M. Gunning and P. Pavlidis. “guilt by association” is not competitive with genetic association for identifying autism risk genes. Scientific Reports, volume 11, Article number: 15950, 2021.

D. M. Gysi, í. D. Valle, M. Zitnik, A. Ameli, X. Gan, O. Varol, H. Sanchez, R. M. Baron, D. Ghiassian, J. Loscalzo, and A. Barabási. Network medicine framework for identifying drug repurposing opportunities for COVID-19. CoRR, abs/2004.07229, 2020. URL https://arxiv.org/abs/2004.07229.

S. Gálvez, A. M. Hirsch, K. L. Wycoff, S. Hunt, D. B. Layzell, A. Kondorosi, and M. Crespi. Oxygen regulation of a nodule-located carbonic anhydrase in alfalfa. Plant physiology, 124(3):1059–68, 2000. ISSN 0032-0889. doi: 10.1104/pp.124.3.1059.

P. E. Hart, N. J. Nilsson, and B. Raphael. A formal basis for the heuristic determination of minimum cost paths. IEEE Trans. Syst. Sci. Cybern., 4(2):100–107, 1968. doi: 10.1109/TSSC.1968.300136. URL https://doi.org/10.1109/TSSC.1968.300136.

Y. Hasin, M. Seldin, and A. Lusis. Multi-omics approaches to disease. Genome biology, 18:1–15, 2017.

K. Hassani-Pak, A. Singh, M. Brandizi, J. Hearnshaw, J. D. Parsons, S. Amberkar, A. L. Phillips, J. H. Doonan, and C. Rawlings. Knetminer: a comprehensive approach for supporting evidence-based gene discovery and complex trait analysis across species. Plant Biotechnology Journal, 19(8):1670–1678, 2021.

Z. He, Y. Luo, X. Zhou, T. Zhu, Y. Lan, and D. Chen. scplantdb: a comprehensive database for exploring cell types and markers of plant cell atlases. Nucleic Acids Research, 52(D1):D1629–D1638, 08 2023. ISSN 0305-1048. doi: 10.1093/nar/gkad706. URL https://doi.org/10.1093/nar/gkad706.

K. Huang, P. Chandak, Q. Wang, S. Havaldar, A. Vaid, J. Leskovec, G. N. Nadkarni, B. S. Glicksberg, N. Gehlenborg, and M. Zitnik. A foundation model for clinician-centered drug repurposing. Nature Medicine, pages 1–13, 2024.

B. Imbert, J. Kreplak, R.-G. Flores, G. Aubert, J. Burstin, and N. Tayeh. Development of a knowledge graph framework to ease and empower translational approaches in plant research: a use-case on grain legumes. Frontiers in Artificial Intelligence, 6:1191122, 2023.

D. Jiao, W. Han, and Y. Ye. Functional association prediction by community profiling. Methods, 129:8–17, 2017.

J. Jumper, R. Evans, A. Pritzel, T. Green, M. Figurnov, O. Ronneberger, K. Tunyasuvunakool, R. Bates, A. Žídek, A. Potapenko, et al. Highly accurate protein structure prediction with alphafold. nature, 596(7873):583–589, 2021.

N. Kavroulakis, E. Flemetakis, G. Aivalakis, and P. Katinakis. Carbon Metabolism in Developing Soybean Root Nodules: The Role of Carbonic Anhydrase. Molecular Plant-Microbe Interactions®, 13(1):14–22, 2000. ISSN 0894-0282. doi: 10.1094/mpmi.2000.13.1.14.

B. Khemani, S. Patil, K. Kotecha, and S. Tanwar. A review of graph neural networks: concepts, architectures, techniques, challenges, datasets, applications, and future directions. Journal of Big Data, 11(1):18, 2024.

D. Kim, J.-G. Joung, K.-A. Sohn, H. Shin, Y. R. Park, M. D. Ritchie, and J. H. Kim. Knowledge boosting: a graph-based integration approach with multi-omics data and genomic knowledge for cancer clinical outcome prediction. Journal of the American Medical Informatics Association, 22(1):109–120, 2015.

J.-Y. Kim, E. Symeonidi, T. Y. Pang, T. Denyer, D. Weidauer, M. Bezrutczyk, M. Miras, N. Zöllner, T. Hartwig, M. M. Wudick, M. Lercher, L.-Q. Chen, M. C. P. Timmermans, and W. B. Frommer. Distinct identities of leaf phloem cells revealed by single cell transcriptomics. The Plant Cell, 33(3):511–530, 01 2021. ISSN 1040-4651. doi: 10.1093/plcell/koaa060. URL https://doi.org/10.1093/plcell/koaa060.

T. N. Kipf and M. Welling. Variational graph auto-encoders. CoRR, abs/1611.07308, 2016. URL http://arxiv.org/abs/1611.07308.

J. Kreplak and J. Burstin. Pisum sativum subsp. sativum cv cameor: Pods, stag…, Nov 2014. URL https://www.ncbi.nlm.nih.gov/biosample/SAMN03164151.

N. Lao and W. W. Cohen. Relational retrieval using a combination of path-constrained random walks. Mach. Learn., 81 (1):53–67, 2010. doi: 10.1007/S10994-010-5205-8. URL https://doi.org/10.1007/s10994-010-5205-8.

P. Larmande and K. Todorov. Agrold: A knowledge graph for the plant sciences. In International Semantic Web Conference, pages 496–510. Springer, 2021.

X. Li, J. Ma, L. Leng, M. Han, M. Li, F. He, and Y. Zhu. Mogcn: a multi-omics integration method based on graph convolutional network for cancer subtype analysis. Frontiers in Genetics, 13:806842, 2022.

X. V. Lin, R. Socher, and C. Xiong. Multi-hop knowledge graph reasoning with reward shaping. In E. Riloff, D. Chiang, J. Hockenmaier, and J. Tsujii, editors, Proceedings of the 2018 Conference on Empirical Methods in Natural Language Processing, Brussels, Belgium, October 31 - November 4, 2018, pages 3243–3253. Association for Computational Linguistics, 2018. doi: 10.18653/V1/D18-1362. URL https://doi.org/10.18653/v1/d18-1362.

Z. Liu, Y. Zhou, J. Guo, J. Li, Z. Tian, Z. Zhu, J. Wang, R. Wu, B. Zhang, Y. Hu, Y. Sun, Y. Shangguan, W. Li, T. Li, Y. Hu, C. Guo, J.-D. Rochaix, Y. Miao, and X. Sun. Global dynamic molecular profiling of stomatal lineage cell development by single-cell rna sequencing. Molecular Plant, 13(8):1178–1193, 2020. ISSN 1674-2052. doi: 10.1016/j.molp.2020.06.010. URL https://www.sciencedirect.com/science/article/pii/S167420522030188X.

W. Ma, Z. Qiu, J. Song, J. Li, Q. Cheng, J. Zhai, and C. Ma. A deep convolutional neural network approach for predicting phenotypes from genotypes. Planta, 248:1307–1318, 2018.

O. Menyhárt and B. Győrffy. Multi-omics approaches in cancer research with applications in tumor subtyping, prognosis, and diagnosis. Computational and structural biotechnology journal, 19:949–960, 2021.

F. Mölder, K. P. Jablonski, B. Letcher, M. B. Hall, C. H. Tomkins-Tinch, V. Sochat, J. Forster, S. Lee, S. O. Twardziok, A. Kanitz, et al. Sustainable data analysis with snakemake. F1000Research, 10:33, 2021.

S. Naithani, P. Gupta, J. Preece, P. D’Eustachio, J. L. Elser, P. Garg, D. A. Dikeman, J. Kiff, J. Cook, A. Olson, S. Wei, M. K. Tello-Ruiz, A. F. Mundo, A. Munoz-Pomer, S. Mohammed, T. Cheng, E. Bolton, I. Papatheodorou, L. Stein, D. Ware, and P. Jaiswal. Plant reactome: a knowledgebase and resource for comparative pathway analysis. Nucleic Acids Research, 48(D1):D1093–D1103, 11 2019. ISSN 0305-1048. doi: 10.1093/nar/gkz996. URL https://doi.org/10.1093/nar/gkz996.

M. Neumann, X. Xu, C. Smaczniak, J. Schumacher, W. Yan, N. Blüthgen, T. Greb, H. Jönsson, J. Traas, K. Kaufmann, et al. A 3d gene expression atlas of the floral meristem based on spatial reconstruction of single nucleus rna sequencing data. Nature Communications, 13(1):2838, 2022.

B. Perozzi, R. Al-Rfou, and S. Skiena. Deepwalk: online learning of social representations. In S. A. Macskassy, C. Perlich, J. Leskovec, W. Wang, and R. Ghani, editors, The 20th ACM SIGKDD International Conference on Knowledge Discovery and Data Mining, KDD ‘14, New York, NY, USA - August 24 - 27, 2014, pages 701–710. ACM, 2014. doi: 10.1145/2623330.2623732. URL https://doi.org/10.1145/2623330.2623732.

O. B. Poirion, Z. Jing, K. Chaudhary, S. Huang, and L. X. Garmire. Deepprog: an ensemble of deep-learning and machine-learning models for prognosis prediction using multi-omics data. Genome medicine, 13:1–15, 2021.

N. Sapoval, A. Aghazadeh, M. G. Nute, D. A. Antunes, A. Balaji, R. Baraniuk, C. Barberan, R. Dannenfelser, C. Dun, M. Edrisi, et al. Current progress and open challenges for applying deep learning across the biosciences. Nature Communications, 13(1):1728, 2022.

M. S. Schlichtkrull, T. N. Kipf, P. Bloem, R. van den Berg, I. Titov, and M. Welling. Modeling relational data with graph convolutional networks. In A. Gangemi, R. Navigli, M. Vidal, P. Hitzler, R. Troncy, L. Hollink, A. Tordai, and M. Alam, editors, The Semantic Web - 15th International Conference, ESWC 2018, Heraklion, Crete, Greece, June 3-7, 2018, Proceedings, volume 10843 of Lecture Notes in Computer Science, pages 593–607. Springer, 2018. doi: 10.1007/978-3-319-93417-4\_38. URL https://doi.org/10.1007/978-3-319-93417-4_38.

R. Schulte-Sasse, S. Budach, D. Hnisz, and A. Marsico. Integration of multiomics data with graph convolutional networks to identify new cancer genes and their associated molecular mechanisms. Nature Machine Intelligence, 3 (6):513–526, 2021.

S. Sehrawat, K. Najafian, and L. Jin. Predicting phenotypes from novel genomic markers using deep learning. Bioinformatics Advances, 3(1):vbad028, 2023.

R. Shahan, C.-W. Hsu, T. M. Nolan, B. J. Cole, I. W. Taylor, L. Greenstreet, S. Zhang, A. Afanassiev, A. H. C. Vlot, G. Schiebinger, P. N. Benfey, and U. Ohler. A single-cell arabidopsis root atlas reveals developmental trajectories in wild-type and cell identity mutants. Developmental Cell, 57(4):543–560.e9, 2022. ISSN 1534-5807. doi: 10.1016/j.devcel.2022.01.008. URL https://www.sciencedirect.com/science/article/pii/S1534580722000338.

Y. Shen, J. Chen, P. Huang, Y. Guo, and J. Gao. M-walk: Learning to walk over graphs using monte carlo tree search. In S. Bengio, H. M. Wallach, H. Larochelle, K. Grauman, N. Cesa-Bianchi, and R. Garnett, editors, Advances in Neural Information Processing Systems 31: Annual Conference on Neural Information Processing Systems 2018, NeurIPS 2018, December 3-8, 2018, Montréal, Canada, pages 6787–6798, 2018. URL https://proceedings.neurips.cc/paper/2018/hash/c6f798b844366ccd65d99bc7f31e0e02-Abstract.html.

J. Stougaard. Regulators and regulation of legume root nodule development. Plant physiology, 124(2):531–540, 2000.

Z. Sun, Z. Deng, J. Nie, and J. Tang. Rotate: Knowledge graph embedding by relational rotation in complex space. In 7th International Conference on Learning Representations, ICLR 2019, New Orleans, LA, USA, May 6-9, 2019. OpenReview.net, 2019. URL https://openreview.net/forum?id=HkgEQnRqYQ.

J. Tang, M. Qu, M. Wang, M. Zhang, J. Yan, and Q. Mei. Line: Large-scale information network embedding. In Proceedings of the 24th international conference on world wide web, pages 1067–1077, 2015.

K. K. Teru, E. G. Denis, and W. L. Hamilton. Inductive relation prediction by subgraph reasoning. In Proceedings of the 37th International Conference on Machine Learning, ICML 2020, 13-18 July 2020, Virtual Event, volume 119 of Proceedings of Machine Learning Research, pages 9448–9457. PMLR, 2020. URL http://proceedings.mlr.press/v119/teru20a.html.

P. Toker, H. Canci, I. Turhan, A. Isci, M. Scherzinger, M. Kordrostami, and E. Yol. The advantages of intercropping to improve productivity in food and forage production–a review. Plant production science, 27(3):155–169, 2024.

T. Trouillon, J. Welbl, S. Riedel, É. Gaussier, and G. Bouchard. Complex embeddings for simple link prediction. In M. Balcan and K. Q. Weinberger, editors, Proceedings of the 33nd International Conference on Machine Learning, ICML 2016, New York City, NY, USA, June 19-24, 2016, volume 48 of JMLR Workshop and Conference Proceedings, pages 2071–2080. JMLR.org, 2016. URL http://proceedings.mlr.press/v48/trouillon16.html.

N. A. Valous, F. Popp, I. Zörnig, D. Jäger, and P. Charoentong. Graph machine learning for integrated multi-omics analysis. British Journal of Cancer, pages 1–7, 2024.

S. Vashishth, S. Sanyal, V. Nitin, and P. P. Talukdar. Composition-based multi-relational graph convolutional networks. In 8th International Conference on Learning Representations, ICLR 2020, Addis Ababa, Ethiopia, April 26-30, 2020. OpenReview.net, 2020. URL https://openreview.net/forum?id=BylA_C4tPr.

L. Wang, J. Liang, Y. Zhou, T. Tian, B. Zhang, and D. Duanmu. Molecular Characterization of Carbonic Anhydrase Genes in Lotus japonicus and Their Potential Roles in Symbiotic Nitrogen Fixation. International Journal of Molecular Sciences, 22(15):7766, 2021a. doi: 10.3390/ijms22157766.

T. Wang, W. Shao, Z. Huang, H. Tang, J. Zhang, Z. Ding, and K. Huang. Mogonet integrates multi-omics data using graph convolutional networks allowing patient classification and biomarker identification. Nature communications, 12(1):3445, 2021b.

Y. Wang, R. Li, D. Li, X. Jia, D. Zhou, J. Li, S. M. Lyi, S. Hou, Y. Huang, L. V. Kochian, and J. Liu. Nip1;2 is a plasma membrane-localized transporter mediating aluminum uptake, translocation, and tolerance in <i>arabidopsis</i>. Proceedings of the National Academy of Sciences, 114(19):5047–5052, 2017. doi: 10.1073/pnas.1618557114. URL https://www.pnas.org/doi/abs/10.1073/pnas.1618557114.

Y. Wang, W. Yu, Y. Cao, Y. Cai, S. M. Lyi, W. Wu, Y. Kang, C. Liang, and J. Liu. An exclusion mechanism is epistatic to an internal detoxification mechanism in aluminum resistance in arabidopsis. BMC plant biology, 20:1–12, 2020.

C. J. Wolfe, I. S. Kohane, and A. J. Butte. Systematic survey reveals general applicability of” guilt-by-association” within gene coexpression networks. BMC bioinformatics, 6:1–10, 2005.

S. Xiao, H. Lin, C. Wang, S. Wang, and J. C. Rajapakse. Graph neural networks with multiple prior knowledge for multi-omics data analysis. IEEE J. Biomed. Health Informatics, 27(9):4591–4600, 2023. doi: 10.1109/JBHI.2023.3284794. URL https://doi.org/10.1109/JBHI.2023.3284794.

B. Yang, W.-t. Yih, X. He, J. Gao, and L. Deng. Embedding entities and relations for learning and inference in knowledge bases. arXiv preprint arXiv:1412.6575, 2014.

B. Yang, W. Yih, X. He, J. Gao, and L. Deng. Embedding entities and relations for learning and inference in knowledge bases. In Y. Bengio and Y. LeCun, editors, 3rd International Conference on Learning Representations, ICLR 2015, San Diego, CA, USA, May 7-9, 2015, Conference Track Proceedings, 2015. URL http://arxiv.org/abs/1412.6575.

S. Zeng, Z. Mao, Y. Ren, D. Wang, D. Xu, and T. Joshi. G2pdeep: a web-based deep-learning framework for quantitative phenotype prediction and discovery of genomic markers. Nucleic acids research, 49(W1):W228–W236, 2021.

D. Zhang, R. Zhao, G. Xian, Y. Kou, and W. Ma. A new model construction based on the knowledge graph for mining elite polyphenotype genes in crops. Frontiers in Plant Science, 15:1361716, 2024.

M. Zhang and Y. Chen. Link prediction based on graph neural networks. In S. Bengio, H. M. Wallach, H. Larochelle, K. Grauman, N. Cesa-Bianchi, and R. Garnett, editors, Advances in Neural Information Processing Systems 31: Annual Conference on Neural Information Processing Systems 2018, NeurIPS 2018, December 3-8, 2018, Montréal, Canada, pages 5171–5181, 2018. URL https://proceedings.neurips.cc/paper/2018/hash/53f0d7c537d99b3824f0f99d62ea2428-Abstract.html.

J. Zhou, G. Cui, S. Hu, Z. Zhang, C. Yang, Z. Liu, L. Wang, C. Li, and M. Sun. Graph neural networks: A review of methods and applications. AI Open, 1:57–81, 2020. doi: 10.1016/J.AIOPEN.2021.01.001. URL https://doi.org/10.1016/j.aiopen.2021.01.001.

X. Zhou, Y. Yi, and G. Jia. Path-rotate: Knowledge graph embedding by relational rotation of path in complex space. In 10th IEEE/CIC International Conference on Communications in China, ICCC 2021, Xiamen, China, July 28-30, 2021, pages 905–910. IEEE, 2021. doi: 10.1109/ICCC52777.2021.9580273. URL https://doi.org/10.1109/ICCC52777.2021.9580273.

Z. Zhu, Z. Zhang, L. A. C. Xhonneux, and J. Tang. Neural bellman-ford networks: A general graph neural network framework for link prediction. In M. Ranzato, A. Beygelzimer, Y. N. Dauphin, P. Liang, and J. W. Vaughan, editors, Advances in Neural Information Processing Systems 34: Annual Conference on Neural Information Processing Systems 2021, NeurIPS 2021, December 6-14, 2021, virtual, pages 29476–29490, 2021. URL https://proceedings.neurips.cc/paper/2021/hash/f6a673f09493afcd8b129a0bcf1cd5bc-Abstract.html.

Z. Zhu, X. Yuan, M. Galkin, L.-P. Xhonneux, M. Zhang, M. Gazeau, and J. Tang. A*net: A scalable path-based reasoning approach for knowledge graphs. In A. Oh, T. Naumann, A. Globerson, K. Saenko, M. Hardt, and S. Levine, editors, Advances in Neural Information Processing Systems, volume 36, pages 59323–59336. Curran Associates, Inc., 2022. URL https://proceedings.neurips.cc/paper_files/paper/2023/file/b9e98316cb72fee82cc1160da5810abc-Paper-Conference.pdf.

